# Nicotine Modifies Responding for Cocaine in a Concurrent Rodent Self-administration Model

**DOI:** 10.1101/2022.09.30.510390

**Authors:** Monica H. Dawes, Paige M. Estave, Stephen E. Albertson, Katherine M. Holleran, Sara R. Jones

## Abstract

Prevailing preclinical models of cocaine use have not resulted in an FDA-approved treatment for cocaine use disorder, potentially due to a focus on cocaine use in isolation, which may not translate well to polysubstance use in clinical populations. Clinically, nicotine has been shown to increase cocaine’s potency and reinforcing efficacy, but some preclinical studies suggest that non-contingent nicotine exposure is not sufficient to alter cocaine self-administration in rats; therefore, this experiment examined if the addition of nicotine to the cocaine solution would alter self-administration behavior. Male Sprague Dawley rats (N=7) were trained to self-administer cocaine (0.75mg/kg/inf), and tested on a long access, fixed ratio 1 schedule of reinforcement (6 hour sessions, unlimited inf, 5 days), for cocaine alone (0.75mg/kg/inf), followed by cocaine and nicotine (0.75mg/kg/inf cocaine+0.03mg/kg/inf nicotine). Finally, rats responded on a progressive ratio schedule for varied doses of cocaine with and without concurrent nicotine at a consistent dose (1.5, 0.75, 0.375, 0.19mg/kg/inf cocaine± 0.03mg/kg/inf nicotine). Unexpectedly, under long access conditions, rats self-administering cocaine and nicotine responded less than for cocaine alone, and did not escalate responding. However, under progressive ratio conditions, responding for cocaine and nicotine was greater than responding for cocaine alone across low and moderate cocaine doses, and decreased at high cocaine doses, indicating a leftward shift in the dose response curve. Together, these data highlight the importance of evaluating multiple outcome measures in nicotine + cocaine paradigms, and suggest that concurrent self-administration of cocaine and nicotine results in greater motivated responding than for cocaine alone.

## 1.1 Introduction

Cocaine use disorder (CUD) is a complex disorder characterized by compulsive drug-seeking, withdrawal, craving, and recurrent use. While there have been more than 300 clinical trials examining potential treatments for CUD, none have resulted in an FDA-approved pharmacotherapy (NIDA, 2020). The identification of medications for CUD frequently begins with preclinical self-administration studies, which have helped to identify potential pharmacological targets. However, a limitation of many preclinical self-administration studies is the examination of a single drug of abuse in isolation, which does not represent the substance use history of clinical populations (Kedia, Sell, & Relyea, 2007; Yiyang Liu, Elliott, Serdarevic et al., 2019; Roy, Richer, Arruda et al., 2013). Self-administration of cocaine alone likely does not encapsulate the complexities of clinical CUD, as people with CUD frequently present with polysubstance use, concurrently using tobacco, alcohol, cannabis, opioids and other drugs (Compton, Valentino, & DuPont, 2021). In particular, nicotine is commonly used in combination with cocaine; there is a two- to four-fold higher prevalence of cigarette smoking in cocaine dependent populations than in the general population (Budney, Higgins, Hughes et al., 1993). Clinical studies of cocaine and nicotine co-use suggest that interactions between these substances may lead to increased consumption of each substance when used in combination (Roll, Higgins, Budney et al., 1996).

Drugs of abuse, including cocaine and nicotine, produce their rewarding effects via the mesolimbic dopamine system, with significant activation of dopaminergic neurons projecting from the ventral tegmental area to the nucleus accumbens (NAc), and interactions between cocaine and nicotine may further increase dopamine release (Pich, Pagliusi, Tessari et al., 1997) (Di Chiara & Imperato, 1988; Pontieri, Tanda, Orzi et al., 1996). In addition, nicotine has been shown to facilitate reward-related dopamine signals by enhancing dopamine during phasic (burst) activity, which may increase reinforced responding (Rice & Cragg, 2004). This suggests that the co-administration of nicotine and cocaine is likely to result in increased responding when compared to cocaine alone.

Results from preclinical and clinical research show enhanced effects following combined cocaine and nicotine use. In clinical populations, concurrent use of cocaine and nicotine results in increased use of both substances (Brewer, Mahoney, Nerumalla et al., 2013; Budney et al., 1993; Kalman, Morissette, & George, 2005; Roll et al., 1996; Sigmon, Tidey, Badger et al., 2003; Wiseman & McMillan, 1996) and enhancement of positive subjective drug effects by self-report measures (Brewer et al., 2013; Wiseman & McMillan, 1998). Preclinically, across animal models and self-administration schedules, acquisition of cocaine self-administration is increased and responding for cocaine is enhanced following noncontingent pretreatment with nicotine (Bechtholt & Mark, 2002; Freeman & Woolverton, 2009; Horger, Giles, & Schenk, 1992; Mello & Newman, 2011; Schwartz, Kearns, & Silberberg, 2018). However, results have been inconclusive regarding the ability of nicotine pretreatment to reinstate previously extinguished responding for cocaine (Bechtholt & Mark, 2002; Schenk & Partridge, 1999), and while nicotine substitutes for the discriminative effects of cocaine in rats, it does not substitute in nonhuman primates, and cocaine does not substitute for nicotine in rodent or nonhuman primate models(Desai, Barber, & Terry, 1999; Mello & Newman, 2011). Additionally, only limited research has been conducted to date regarding concurrent, contingent nicotine and cocaine self-administration in animal models (Barbosa-Mendez & Salazar-Juarez, 2018; Freeman & Woolverton, 2009; Mello, Fivel, & Kohut, 2013; Mello, Fivel, Kohut et al., 2014; Mello & Newman, 2011).

It is well-known that drug-associated cues play a large role in the formation and continuation of drug taking behaviors, and the presence of drug-paired cues is of particular importance in nicotine self-administration behavior (Butler, Forget, Heishman et al., 2021; Caggiula, Donny, Palmatier et al., 2009; Caggiula, Donny, White et al., 2001; Chaudhri, Caggiula, Donny et al., 2007; Donny, Caggiula, Knopf et al., 1995; Donny, Chaudhri, Caggiula et al., 2003; LeSage, Burroughs, Dufek et al., 2004; X. Liu, Caggiula, Yee et al., 2006; Sorge, Pierre, & Clarke, 2009). Nicotine is also thought to enhance the salience of cues and support the development of habitual behaviors (Asgaard, Gilbert, Malpass et al., 2010; Claus, Blaine, Filbey et al., 2013; Lê, Wang, Harding et al., 2003; Overby, Daniels, Del Franco et al., 2018), which may play a role in increasing cocaine self-administration. As such, the present study seeks to examine the behavioral consequences of combined cocaine and nicotine relative to cocaine alone across several behavioral paradigms. Results from this study suggest that concurrent self-administration of cocaine and nicotine increases the reinforcing efficacy of cocaine, with decreased long access responding and increased progressive ratio responding.

## 1.2 Materials and Methods

### Animals

Male Sprague-Dawley rats (350-400 g; Envigo, Indianapolis, IN) were maintained on a 12:12 hour reverse light/dark cycle (0300 off; 1500 on) with *ad libitum* food and water. All animals were maintained according to the National Institutes of Health guidelines in Association for Assessment and Accreditation of Laboratory Animal Care accredited facilities. The experimental protocol was approved by the Institutional Animal Care and Use Committee at Wake Forest School of Medicine.

### Drugs

Cocaine HCl was acquired from the National Institute on Drug Abuse Drug Supply Program (Bethesda, MD) dissolved in sterile saline. (-)-Nicotine hydrogen tartrate salt (Cayman Chemical, Ann Arbor, MI) was dissolved in sterile saline (pH adjusted to 7.4).

### Apparatus

Rats were housed and trained in custom-made operant conditioning chambers located within the standard housing room. In all sessions, an active lever response resulted in retraction of the lever and illumination of the stimulus light during the 20s timeout. All sessions were completed during the active dark cycle (0900 to 1500).

### Food Training

One week after arrival, rats were individually housed and trained to self-administer sucrose pellets (45mg) on an FR1 schedule (4s timeout, 75 pellet maximum, 6 hr sessions). Once a minimum of 50 pellets were earned with at least a 2:1 preference for the active over inactive lever, food training was stopped. Rats that did not meet the criteria within five sessions were removed from the study (Exclusion N=1).

### Self-Administration Surgery and Training

Rats were anesthetized before implantation with a chronic indwelling jugular catheter as described previously (Y. Liu, Roberts, & Morgan, 2005). After a two day recovery period, animals were trained to self-administer cocaine on an FR1 schedule of reinforcement. Each lever press resulted in the intravenous delivery of 0.75mg/kg/inf cocaine over four seconds. Rats could earn up to 20 infusions per 6 hr session. Acquisition occurred when an animal responded for 20 injections for two consecutive days. Experimental timeline and days to acquisition are shown in Figures 1A and 1B, respectively.

**Fig. 1.**
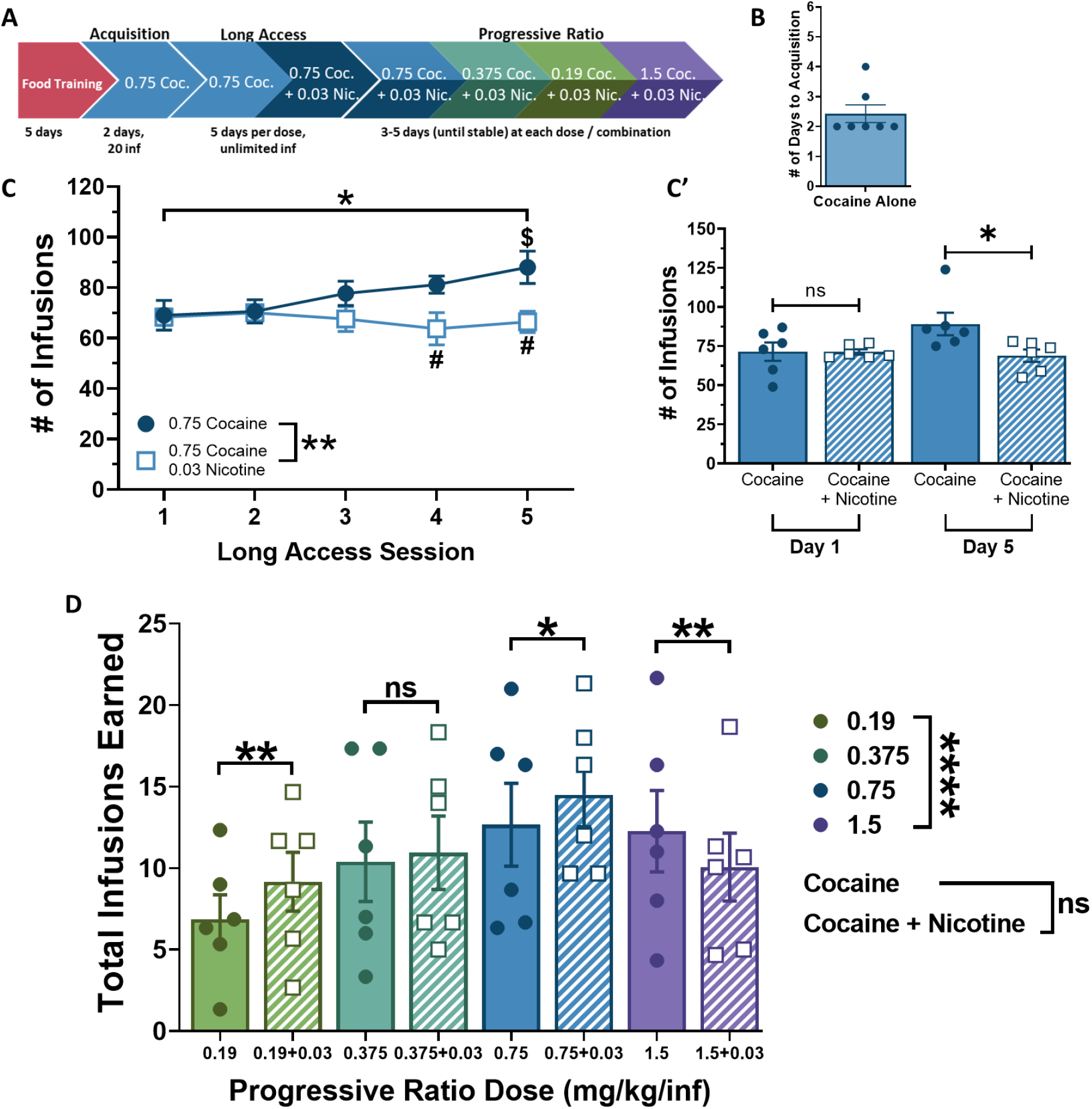
*A)* Experimental timeline. Order of presentation of cocaine alone or cocaine and nicotine combination during progressive ratio responding was randomized for each rat. *B)* Mean (± SEM) number of sessions required to reach acquisition criteria, 2.429 days ± 0.297, N=7. *C)* Self-administration responding across five days of long access, cocaine alone and cocaine + nicotine, N = 6. Significant main effects of session and nicotine condition, significant session X nicotine condition interaction. (#) significant difference between nicotine conditions ($) significant difference from session 1. *C’)* Comparison of self-administration responding on days 1 and 5 between nicotine exposure conditions, N=6. No significant difference between conditions on day 1 of exposure, significant difference on day 5 of exposure. *D)* Progressive ratio dose response curves for cocaine alone and cocaine + nicotine, N = 6. Significant main effect of cocaine dose, no significant effect of nicotine condition, significant cocaine dose X nicotine condition interaction. Post-hoc testing indicated significant differences between nicotine conditions at 0.19, 0.75, and 1.5 mg/kg/inf cocaine doses.

### Long Access

Rats were allowed unlimited access (FR1) to 0.75mg/kg/inf cocaine during 6 h sessions for five consecutive days. Next, all animals were allowed unlimited access (FR1) to the combination of 0.75mg/kg/inf cocaine + 0.03mg/kg/inf nicotine (mixed in the same syringe) under the same conditions for a further five days.

### Progressive Ratio

Rats self-administered on a progressive ratio schedule as described previously (Karkhanis, Beveridge, Blough et al., 2016). The outcome measure used was total number of infusions. Cocaine doses were administered in decreasing order (0.75, 0.375, 0.19mg/kg/inf) then increased to 1.5mg/kg/inf. Initial exposure condition (± nicotine) was randomized for each rat at each cocaine dose. Exposure conditions were changed once stable responding (three days with ≤20% change in total number of infusions earned) or five total days was reached. Animals that did not stabilize on any dose were excluded (Exclusion N=1).

### Statistics

All statistical analyses were conducted using GraphPad Prism 8 (Graph Pad Software, La Jolla, CA). Data are presented as mean ± SEM, significance level p<0.05. Repeated measure two-way ANOVAs were used to determine significance and were followed by Sidak’s post-hoc tests as appropriate.

## 1.3 Results

### Concurrent cocaine + nicotine decreases responding for cocaine and prevents escalation of cocaine-taking behaviors

Significant main effects of session (F_2,24_=2.957, p=0.0405) and nicotine presence (F_1,6_=27.26, p=0.002) were observed across the five days of long access self-administration (Figure 1C), with a significant interaction of session X nicotine condition (F4,24=5.089, p=0.0041). Sidak post-hoc tests revealed a significant increase in responding from session 1 to session 5 of cocaine alone self-administration (t_(7, 24)_=4.492, p=0.0015), but no significant change in responding from session 1 to session 5 of cocaine + nicotine self-administration (t_(7, 24)_=0.4390, p>0.9999). Responding on day 1 of concurrent cocaine + nicotine was not significantly different than responding on day 1 of cocaine alone (t_(7, 24)_=0.1689, p>0.9999; Figure 1C’), but responding on days 4 & 5 of concurrent cocaine + nicotine was significantly lower than responding on day 5 of cocaine alone self-administration (t_(7, 24)_=5.100, p=0.0002; Figure 1C’).

### Concurrent nicotine alters responding for cocaine on a progressive ratio schedule of reinforcement

A significant main effect of cocaine dose was observed across the progressive ratio dose response curve (F_3,42_=211.69, p<0.0001), with a significant interaction of cocaine dose X nicotine condition (F_3,42_=10.00, p<0.001), but no significant main effect of nicotine condition (F_1,14_=0.6711, p=0.4264; Figure 1D). Post hoc analysis determined that, at lower doses of cocaine (0.19 – 0.75 mg/kg/inf), the addition of nicotine increased responding (0.19—t_(42)_=3.434, p=0.0054; 0.375— t_(42)_=0, p>0.9999; 0.75—t_(42)_=2.814, p=0.0294), while at the highest dose of cocaine (1.5 mg/kg/inf) the addition of nicotine significantly decreased responding (t_(42)_=3.492, p=0.0046).

## 1.4 Discussion & Conclusions

This study presents a novel examination of concurrent intravenous cocaine and nicotine self-administration under long access and progressive ratio schedules of reinforcement. Concurrent nicotine reduced responding for cocaine on an FR1 schedule and at the highest cocaine dose of the PR dose response curve. In contrast, nicotine increased responding for cocaine at the lowest and intermediate cocaine doses during the PR dose response curve. Based on these results, we suggest that nicotine acts to enhance cocaine’s rewarding efficacy, requiring fewer infusions to reach equivalent reward values.

Results from clinical CUD research suggest that nicotine enhances cocaine intake (Budney et al., 1993; Roll, Higgins, & Tidey, 1997; Wiseman & McMillan, 1996), and results in enhanced positive subjective effects of cocaine use (Wiseman & McMillan, 1998) when compared to cocaine use alone. Preclinical findings mirror clinical results, as multiple animal models of self-administration show faster acquisition of cocaine taking and greater responding for cocaine across varied reinforcement schedules, following noncontingent pretreatment with nicotine (Bechtholt & Mark, 2002; Horger et al., 1992; Mello & Newman, 2011). Despite evidence suggesting that nicotine alone does not produce robust reinforcing effects (Caggiula et al., 2009; Dougherty, Miller, Todd et al., 1981), it has been shown to facilitate cocaine reinforcement, as repeated pretreatment with nicotine produces leftward shifts in cocaine PR dose response curves (Bechtholt & Mark, 2002).

Nicotine increases mesolimbic dopamine release via nicotinic acetylcholine receptors (nAChRs), and there is evidence that suggests cocaine acts as a nAChR antagonist (Acevedo-Rodriguez, Zhang, Zhou et al., 2014; Chen, Gao, Ma et al., 2019). Therefore nAChRs may represent a target of interest for future studies of combined cocaine and nicotine self-adminsitration.

As this study is one of the first to examine concurrent, contingent self-administration of combined cocaine and nicotine, there are some potential limitations of this paradigm, and complexities of comparing these findings to previous literature. First, this study did not replicate previous findings using noncontingent nicotine administration by Bechtholt & Mark (2002), which showed escalation of cocaine self-administration behavior following repeated nicotine pretreatment. As the present study used five days of long access self-administration compared to Bechtholt & Marks’ 14 days, there may not have been sufficient nicotine exposure to replicate the long-term effects seen previously (Bechtholt & Mark, 2002). However, as the Bechtholt & Mark study used noncontingent nicotine pretreatment prior to cocaine self-administration, and the study discussed here uses contingent nicotine in concurrence with cocaine, it is likely that methodological considerations account for the difference in results, rather than insufficient nicotine exposure. The dose of nicotine used in this experiment (0.03mg/kg/inf) has been used extensively in nicotine alone self-administration (Donny et al., 1995; LeSage et al., 2004; Sorge et al., 2009), and is therefore unlikely to be aversive. To avoid the development of tolerance to cocaine which may occur following repeated self-administration of high doses of cocaine, doses were presented in descending order (0.75, 0.375, 0.19mg/kg/inf), with the highest dose of cocaine presented last (1.5mg/kg/inf); however, this could be a limitation of this study, as all possible dose orders were not examined.

An important methodological consideration of this experiment are the access schedules used, and the drug-associated cues related to cocaine and nicotine self-administration. Drug-paired cues are critical for nicotine self-administration, and there are notable differences in the methodology used in this experiment compared to standard nicotine self-administration methods. Nicotine self-administration is typically completed in short, limited access sessions of one to two hours (Perkins, 1999), and is highly influenced by environmental and contextual cues, such as visual signals or the separation of housing and drug-associated chambers (Caggiula et al., 2001). In this experiment, rats lived in the self-administration chambers, sessions lasted six hours, and cocaine + nicotine were administered concurrently through the intravenous catheter following each active lever response. Together these factors reduced the number of nicotine-associated cues and potentially decreased responding for combined cocaine + nicotine. Additionally, while this model has greater translational validity compared to cocaine alone self-administration models, while humans use cocaine and nicotine at the same time, the routes of administration differ, while the rats in this study received their cocaine and nicotine concurrently following each lever press. Future studies implementing a cocaine vs. nicotine choice procedure may provide further insight into cocaine and nicotine polysubstance use.

In conclusion, concurrent cocaine and nicotine infusions in rats trained to self-administer cocaine reduced responding on an FR1 long access schedule and produced left-wards shifts in a PR dose response curve, suggesting an increase in cocaine’s reinforcing effects when administered concurrently with nicotine. While in human populations, nicotine may act as a cue for polysubstance use when cocaine is available, our study supports that nicotine itself, without environmental context cues, alters the rewarding value of cocaine. This information is relevant for treatment development for CUD and suggests future strategies may need to consider nicotine use in conjunction with CUD, as the data presented here show augmentation of cocaine taking behaviors by nicotine, potentially due to increased cocaine potency.

